# Characterising the periodontal granulation tissue using scRNAseq

**DOI:** 10.1101/2025.01.13.632112

**Authors:** Wentao Zhu, Kathy Fung, Pawan Dhami, Paul Sharpe, Jan Krivanek, Luigi Nibali, Cheng Zhang, Vitor C.M. Neves

## Abstract

**Aims:** Our research investigated the cellular composition and molecular mechanisms of periodontal granulation tissue using single-cell RNA sequencing (scRNA-seq), aiming to enhance the understanding of periodontal disease pathogenesis and identify potential targets for regenerative therapies.

**Materials and Methods:** Granulation tissue samples were collected from patients undergoing periodontal surgery. Fresh tissues were processed into single-cell suspensions and subjected to scRNA-seq. The data were integrated with existing datasets from healthy gingiva and periodontal ligament. Computational analyses were performed, and validation through immunofluorescence staining.

**Results:** Ten distinct cell clusters were identified across the samples. Granulation tissue exhibited a higher abundance of immune cells compared to healthy tissues. A novel endothelial cell subpopulation, exclusive to granulation tissue, was discovered, characterized by NOTCH3 expression and involvement in ossification pathways. Additionally, granulation tissue Fibroblast subpopulations demonstrated a progenitor-like state, characterized by extracellular matrix reorganization and low differentiation, similar to cancer-associated fibroblasts.

**Conclusion(s):** This study advances the understanding of periodontal disease by characterizing key regulatory cell populations within granulation tissue. The identification of the novel endothelial subpopulation offers new insights into the disease’s pathogenesis and presents potential targets for regenerative therapies. These findings suggest opportunities for developing biomaterials that modulate specific cellular pathways to improve periodontal disease treatment.

## Introduction

The gingiva, characterized by a keratinized stratified squamous epithelium and an underlying connective tissue matrix, is an actively proliferating tissue, serving as the frontline defence against oral diseases. This tissue is responsible to maintain biological equilibrium amidst various microenvironmental challenges, however, inadequate oral hygiene practices, facilitating the accumulation and calcification of plaque—a bacterial biofilm—can disrupt this delicate balance[1]. Consequently, host immune responses precipitate dysbiosis within the oral microbiota leading to periodontal disease, a spectrum of inflammatory conditions afflicting the periodontal tissues, culminating in irreversible damage to the supporting structures of the teeth[2].

The sequelae of periodontal disease manifest as the destruction of attachment, including bone and ligaments, accompanied by a transformation of gingival tissue from its anatomically defined state to a fibrous and inflamed granulation tissue reminiscent of scar formation[3]. Traditionally, granulation tissue has been excised during periodontal therapy[4]. However, emerging evidence suggests that this tissue may harbour cells with regenerative and ossification potential, thus prompting reconsideration of its therapeutic implications[5].

Recent clinical observations and case-based research suggest that whether or not to remove granulation tissue is questionable, as the cellular composition and contributions to the formation of granulation tissue remains poorly understood[6]. This knowledge gap underscores the challenges associated with regenerating the periodontium in patients with periodontitis, making the process both challenging and unpredictable. The characterisation of the granulation tissue cell component and its origins can result in profound implications for understanding the pathogenesis of periodontal disease and may provide new avenues for regenerative therapies.

Therefore, this research used single cells RNA sequencing (scRNAseq) to investigate the intricate composition of the granulation tissue and elucidate its cellular underpinnings. Our high-resolution findings have identified similar potential pathways of differentiation from health to disease state, as well as a new endothelial cell population exclusive of the granulation tissue with higher potential of ossification. Together our data advances our understanding of periodontal disease and ultimately shine a light onto novel targets to improve regenerative strategies and improvement of periodontal disease treatment.

## Materials and methods

Patients undergoing routine periodontal surgical procedures at the Department of Periodontology, Guy’s Hospital, King’s College London, provided consent for the collection of human granulation tissue samples. The study protocol adhered to the guidelines set forth by the UK Human Tissue Act and received ethical approval from the East of England-Cambridge East Research Ethics Committee (reference 20/EE/0241). Prior to inclusion in the study, written informed consent was obtained from all participants. Inclusion criteria for the patient cohort stipulated the absence of relevant medical conditions, non-usage of prescribed medication, non-usage of nicotine or nicotine-replacement products, and non-pregnancy or breastfeeding status. The granulation samples originated sites indicated for surgical intervention post provision of step 1 and step 2 of periodontal treatment according the S3 guidelines and samples included in the sequencing are referent from the following surgical sites:

Patient 1 - UR1 intrabony granulation (Male – 59 years old – fit and well)

Patient 2 - UR6 supracrestal granulation (Female – 37 years old – fit and well)

Patient 3 – LL6 intrabony and supracrestal granulation (Female – 30 years old – fit and well)

Fresh tissue obtained during surgery from intrabony and suprabony defects were dissected and dissociated into a single cell suspension. Following mechanical dissociation with scalpel, the samples were placed in PBS + 2% FBS with 2 mg/ml of Collagenase Type II and 1 mg/ml of DNAse Type I (Merck), and incubated for 20–30⍰min at 37⍰°C shaking (120⍰rpm) with homogenization of the suspension every 3-4 minutes. After incubation, the cell suspensions were toped up to 12ml using using PBS + 2% FBS and filtered in using a 40μm cell strainer. These were subsequently centrifuged in 4⍰°C pre cooled centrifuge for 5⍰min at 300⍰× ⍰*g*. The samples were then cleansed for debris using Debris Removal Solution (Miltenyi Biotec) according to the manufacturer. The pellet was then re-suspended in 100 uL of PBS + Ultrapure™BSA (0.04%) (Thermo-Fischer Scientific) and sent to the BRC Genomic Centre where the library preparation and sequencing were performed.

Single-cell suspensions were manually counted using a haemocytometer and concentration adjusted to a minimum of 300cells/μL. Cells were loaded according to standard protocol of the Chromium single-cell 3’ kit to capture around 5,000 cells per chip position. Briefly, a single-cell suspension in PBS 0.04% BSA was mixed with RT-PCR master mix and loaded together with Single Cell 3’ Gel Beads and Partitioning Oil into a Single Cell 3’ Chip (10x Genomics) according to the manufacturer’s instructions. RNA transcripts from single cells were uniquely barcoded and reverse transcribed. Samples were run on individual lanes of the Illumina HiSeq 2500.

### Data Pre-processing and Quality Control

Raw sequencing data from granulation tissue, along with datasets from GSE152042 (gingiva) and GSE161267 (periodontal ligament), were demultiplexed and mapped to the human reference genome (GRCh38) using Cell Ranger 6 (v5.0)[7]. Gene expression matrices were generated and processed for quality control in RStudio (v4.1.2) using Seurat (v5.1.0)[8]. During quality control, genes expressed in fewer than three cells were removed. Cells with more than 8,000 unique feature counts or fewer than 600 unique feature counts were excluded to filter out potential doublets, poor-quality cells, or empty droplets. Additionally, cells with mitochondrial gene expression exceeding 25% were removed to ensure data integrity. After pre-processing, the dataset consisted of 6,729 cells from disease granulation tissue (‘D’), 4,001 cells from gingiva (‘G’), and 3,024 cells from periodontal ligament (‘L’).

### Computational Analysis of Integrated scRNA-seq Datasets

Based on the filtered cells, data normalization was first performed to adjust for sequencing depth. The 2,000 most variable features in each dataset were identified using the variance-stabilizing transformation method. Next, integration anchors were computed across the datasets, and batch effects were removed through data integration. This integration approach was carefully compared with a non-integration strategy to assess alignment with known subcluster characteristics. Differences between clusters were attributed to tissue-specific gene expression variations rather than batch effects, based on the subcluster analysis, thereby ensuring the biological accuracy of the findings.

Following these steps, the data were scaled to account for sequencing depth and other technical variations. Principal component analysis (PCA) was performed to identify key dimensions for clustering. A k-nearest neighbor (KNN) graph was constructed to establish relationships between cells, and clustering was carried out using the Louvain algorithm, optimizing modularity with a resolution parameter of 0.5. To visualize cell distribution, uniform manifold approximation and projection (UMAP) was applied, focusing on the first 30 principal components and incorporating 30 nearest neighbors. Marker genes were then identified by analyzing differential expression using the non-parametric Wilcoxon rank sum test, as implemented in Seurat’s default FindMarkers function. Cell type annotation was performed based on the expression of specific marker genes within each cluster. Cell types identified from our datasets included plasma cells (MZB1), endothelial cells (EMCN, CLDN5), T cells (TRAC, TRBC1), epithelial cells (KRT15, KRT14), fibroblasts (COL1A1, COL6A1, FAP), mesenchymal stem cells (RGS5, ACTA2, FRZB), macrophages (LYZ), memory B cells (BANK1), and mast cells (HDC). The labeled UMAP was visualized either on the merged dataset or on each of the three tissues individually. Additionally, the percentage of each cell type was calculated and clustered across the tissues.

### Subcluster re-clustering

Endothelial, epithelial, fibroblast, and mesenchymal stem cell populations were identified within their respective clusters in the UMAP plot and were subsequently re-clustered following the same protocol outlined above for the second round of analysis. For this re-clustering, UMAP was utilized with a resolution parameter set to 0.25.

### Gene Ontology (GO) analysis

The ClusterProfiler package in R [9] was used for functional enrichment analysis using the molecular functions and biological pathways gene ontology (GO) annotations and Kyoto Encyclopedia of Genes and Genomes (KEGG) pathways. The enrichment addressed the marker list of each cluster.

### Cell-Cell Communication

Cell-cell communication within the subclusters was analyzed using the CellChat pipeline (v1.1.3) [10]. A new CellChat object was created from the merged Seurat object, and the complete CellChat database, including ECM signaling, cell-cell contact, and secreted signaling pathways, was selected for analysis. Communication probabilities were computed using a truncated mean approach (computeCommunProb function, type = “truncatedMean”, trim = 0.2). Subsequently, a cell-cell communication network was inferred, and downstream visualization techniques were applied to depict the interactions.

### Single-cell trajectory analysis

Pseudotime analysis was performed using Monocle2 [11] with the DDR-Tree algorithm and default parameters, marker genes for each subcluster were selected, and raw expression counts from cells that passed quality control filtering were utilized, then, the analysis was conducted to model the differentiation trajectories, and Branch Expression Analysis Modeling (BEAM) was applied to identify genes associated with branch-specific fate decisions. The starting point of the pseudotime trajectory was determined using CytoTRACE2 [12], based on the differentiation potential predicted by the model. This integrated approach enabled the assessment of differentiation levels among clusters and validated the pseudotime starting point by integrating marker genes that exhibited differential expression according to our list of marker genes.

### Gene Set Enrichment Analysis (GSEA)

Differentially expressed genes that induced functional pathways between endothelial cells from different sources were compared, with contrasts made between “D” versus “L” and “D” versus “G”. Gene Set Enrichment Analysis (GSEA, v4.0.3) [13] was performed to evaluate endothelial cell-induced vascular development between granulation and healthy tissues. The top 5 enriched pathways were visualized using a dot plot, with rankings based on the enrichment P-value.

### Histology

Granulation samples collected were fixed in 4% paraformaldehyde (PFA) for 24 hours at 4°C. The samples were processed and mounted in O.C.T. and cut in the cryostat (BRIGHT, OTF5000) at 8μm thickness. To visualise the tissue morphology, the slides were stained using Haematoxylin & Eosin staining and viewed in a light-field Nikon Eclipse Ci-L microscope.

### Immunofluorescence

To visualise and validate the findings of the sequencing, the samples were stained with mouse polyclonal anti-VWF antibody (1:100; Abcam; ab201336), rabbit polyclonal anti-RGS5 antibody (1:50; Thermofisher; 11590-1-AP), rabbit polyclonal anti-NOTCH3 antibody (1:250; Abcam; ab23426), rabbit polyclonal anti-HEY1 antibody (1:50; Abcam; ab154077) overnight at 4 °C. Sections were washed and exposed to secondary antibody (1:250; Thermo Fisher Scientific; A21449) for 1 h at room temperature. Imaging was generated using a Thunder microscope.

## Results

### Generation of the comprehensive transcriptional landscape in disease-progressing granulation tissue

In this research, we aimed to unravel the intricate transcriptional landscape underlying periodontal inflammation by conducting scRNA-seq analysis of the diseased granulation periodontal tissue, comparing to healthy gingiva, and periodontal ligament. Our findings shed light on the molecular complexity of these tissues and provide valuable insights into the pathogenesis progression of periodontal disease. A comprehensive experimental workflow was designed, detailing the steps involved in sample collection, processing, sequencing, and data analysis for the characterisation of the granulation tissues (Supplementary Figure 1). This approach allowed us to obtain high-resolution transcriptomic data from a diverse range of cell populations present in the periodontal microenvironment. Starting with an overview of the scRNA-seq analysis, we integrated our dataset with previously publicly available datasets GSE152042 (gingiva)[14] and GSE161267 (periodontal ligament)[15]. By applying stringent quality filtering methods, we obtained single-cell transcriptomes from a total of 13,754 single cells (granulation tissue = 6,729, gingiva = 4,001, periodontal ligament = 3,024). Unbiased clustering of the cells identified ten clusters based on uniform manifold approximation and projection (UMAP) analyses (Figure 1A,B), and the cell types were annotated (Figure 1C,D) according to previously published datasets [14,15]. We investigated the proportion of each cell cluster comparing between the sample sets (Figure 1E,F). Our results identified that diseased granulation tissue had a higher abundance of immune/plasma cells (69.5% of the total cell count) compared to healthy gingiva and periodontal ligament (23.7% and 14.8% respectively). KEGG pathway enrichment analysis showed that the immune compartment was enriched for osteoclast differentiation pathways (Supplementary Figure 2). We also identified a small population of epithelial cells (3.3%), mesenchymal stem cells (5.5%), and endothelial cells (11%) in the granulation tissues. Interestingly the fibroblast population proportion was not significantly different from the healthy tissues. These findings deepen our understanding of the cellular composition and molecular signatures associated with periodontal granulation tissues.

**Figure 1.**
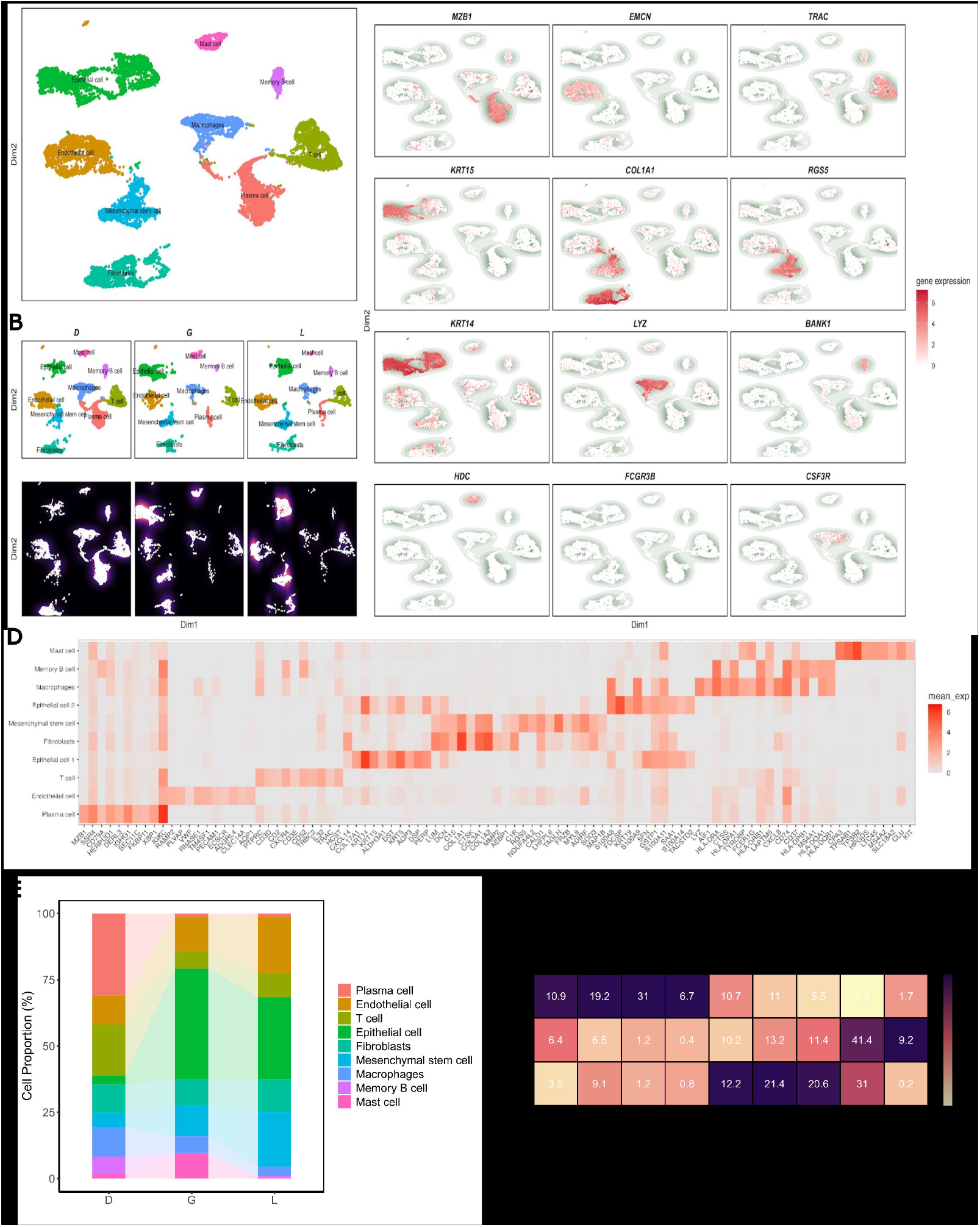
Characterising the populations of periodontal tissues. (A) Uniform Manifold Approximation and Projection (UMAP) representation showing nine cell types identified in the granulation tissue. (B) UMAP of the *data obtained from the separate tissues involved in analysis D refers to Diseased (Granulation tissue), G refers to Gingiva (Healthy), and L refers to Periodontal Ligament (Healthy) (Cell number per tissue: Diseased granulation tissue = 6,729, Gingiva = 4,001, Periodontal Ligament = 3,024)*. (C) Selected markers used for cell type identification correlating to the UMAP. (D) Differentially expressed genes (DEGs) were normalized and visualized on a heatmap, displaying the Top 10 genes ranked by log2 fold change (log2FC). (E,F) Rate of cell annotation from each tissue resource.

### *In silico* dissection of the granulation tissue

To dive into the role of the different cell compartments in the granulation tissue, we performed UMAP subcluster analysis. We first conducted a cell-cell communication analysis by using the CellChat package, on the collective of these three tissues to understand which populations we should focus on. Our results demonstrate that the populations producing the biggest weight of interactions were the Fibroblasts, Endothelial cells, Epithelial cells, and Mesenchymal stem cells (Supplementary Figure 3A), therefore our subcluster analysis focused on these populations. Our cell-cell communication analysis also highlighted that both endothelial and fibroblast cells clusters exhibit the most pronounced changes in signaling pathways (Supplementary Figure 3B). Specifically, endothelial cells are shown to receive signals, while fibroblasts are responsible for their release. This interaction underscores the critical roles these two cell types play in mediating communication and functional responses within the tissue microenvironment.

### Subcluster analysis of the Epithelial cells

The proportion of epithelial cells in granulation tissue is significantly lower compared to the gingiva and ligament (Figure 1E). Inside of this small epithelial compartment, a total of 5 subcluster were identified in which two subclusters were of mixed origin (M_EP3 and M_EP4) and the other three were specific of each tissue, granulation (D_EP), gingiva (G_EP1), and PDL (L_EP1) (Supplementary Figure 4). Gene ontology (GO) analysis of the populations revealed distinct enrichment for each subpopulation with D_EP enriching for wound healing and regulation of cell projection adhesion. This subset showed upregulation of differentiation markers, such as KRT13 and CLDN1, alongside the downregulation of basal markers KRT14 and KRT15, indicating a transition from progenitor activity toward differentiation to restore the junctional epithelial barrier (Supplementary Figure 5)[16]. Additionally, D_EP subpopulation expresses higher levels of VIM indicating a potential epithelial-mesenchymal transition (EMT), which may facilitate bacterial invasion into the underlying gingival tissues and propagation of inflammation[17]. To understand the potency of differentiation capacity of the epithelial populations we have employed Pseudotime and CytoTRACE2 analyses on these populations revealing that D_EP is highly differentiated in comparison to the G_EP1, suggesting that these epithelial cells have low developmental potential.

### Subcluster analysis of the Mesenchymal stem cells

Within the MSC compartment, 6 subclusters were identified, of which one subcluster was specific to granulation tissue (D_MSC) (Supplementary Figure 6). GO analysis of the populations revealed that D_MSC was enriched for small gtpase mediated signal transduction and amebiodal-type cell migration, suggesting that this subcluster plays a pivotal role in dynamic cellular processes involved in tissue remodeling and repair. The enrichment for small GTPase-mediated signal transduction indicates a regulatory function in cytoskeletal organization[18], while the association with amoeboid-type cell migration suggests a capacity for rapid and adaptive movement through the extracellular matrix[19], facilitating granulation tissue formation. These findings highlight the specialized functions of D_MSC within the granulation tissue microenvironment. Pseudotime and CytoTRACE2 analyses revealed that D_MSC is highly differentiated in comparison to PDL and Gingival MSC populations, with the cells from the G_MSC1 population comprising the majority on the trajectory of becoming D_MSC, and a small proportion of L_MSC1 on the same trajectory. Altogether, these findings suggested that a small subpopulation of gingival and PDL cells take on a differentiation commitment into the D_MSC population, and that the D_MSC population displays a distinct functional trajectory to the Gingiva and PDL MSCs.

### Subcluster analysis of the Fibroblasts

The fibroblast subclustering analysis revealed 4 subclusters, with the granulation subcluster (D_FB1) being enriched in the functional GO analysis for extracellular matrix organization, extracellular structure organization, and external encapsulating structure organization, suggesting that D_FB1 plays a critical role in shaping the extracellular matrix (ECM) architecture during granulation tissue formation (Figure 2A-G). Cell-to-cell interaction analysis of the fibroblast subcluster revealed that D_FB1 had the biggest weight of interaction of outgoing and ingoing signaling patterns, singling and receiving Collagen, Laminin, Midkine, and Fibronectin, all of which could be therapeutic targets (Figure 2H,I). Pseudotime and CytoTRACE2 analyses revealed that D_FB1 is highly undifferentiated in comparison to PDL and Gingival MSC populations (Figure 2J), suggesting that D_FB1 represents a progenitor-like fibroblast population that is poised to respond dynamically to reparative cues in the granulation tissue microenvironment. Additionally, the granulation tissue also shows a slight but specific expression of LEF1 (Supplementary Figure 7), a key transcription factor in the Wnt signaling pathway that regulates cell proliferation and differentiation; and of increased expression of HSPA1A and HSPA1B, which are genes associated with metastatic colon cancer [20] (Supplementary FIgure 8).This supports the idea that the fibroblasts in this tissue are more inclined towards a low-differentiation state. This unique profile indicates that D_FB1 could be the critical factor for the scar-like formation phenotype seen in the granulation tissue.

**Figure 2.**
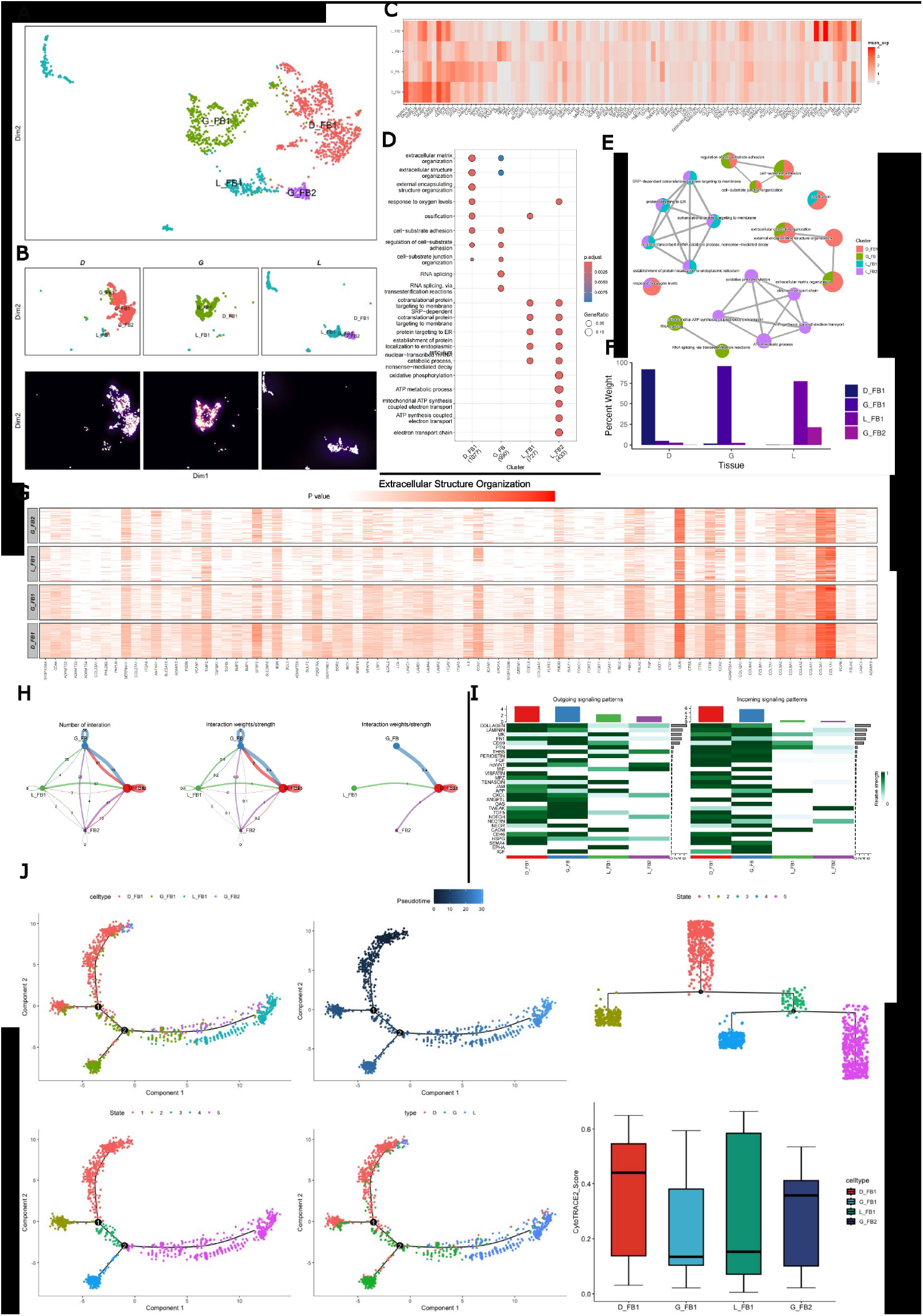
Fibroblast subpopulation analysis. (A) Reclustering UMAP plot of Fibroblast cells, single cells coloured by cluster annotation. (B) UMAP plot divided by tissue and population density plot. (C) Differentially expressed genes (DEGs) were normalized and visualized on a heatmap, displaying the Top 10 genes ranked by log2 fold change (log2FC). (D,E) GO enrichment terms for comparing clusters, using the compareCluster function, and emapplot function from clusterProfiler. (F) Rate of cell annotation from each subpopulation. (G) Extracelular Structure Organisation genes and their expression levels were plotted on a heatmap. (H) Circos plot depicts cell interactions among the four cell sub-types within the Fibroblast populations. Node size corresponds to the number of interactions, while edge width indicates the number of significant ligand-receptor pairs between two cell types. (I) Signaling sender and receiver information and enriched as clusters. (J) Pseudotime plotting reveals the differentiation trajectory of Fibroblasts cells, annotated by cell type, state, cluster, and tissue source, and CytoTrace2 identification of the starting point of fibroblast cell differentiation.

### Subcluster analysis of the Endothelial cells

Finally, to identify the role of the endothelial cells in the periodontal tissues, sub cluster analysis revealed the presence of five different subclusters, amongst which two were granulation tissue specific (V_EC1 and V_EC4) (Figure 3A,B). To characterise the vascular endothelium, we created a marker gene plot which showed that Von Willebrand Factor (VWF) was identified in all clusters establishing its identity as a vascular endothelial marker (V_EC)(Figure 3C). Conversely, LYVE1, indicative of lymphatic endothelium[21], was exclusively detected in cluster 5 (L_EC), originating mainly from gingiva. GO enrichment analyses revealed a pronounced involvement of ossification pathways was observed in endothelial cells (V_EC1 and V_EC4), particularly in cluster V_EC4, which originates specifically from granulation tissue (Figure 3D,E). To validate the presence of the V_EC4 subcluster in the granulation tissue, we identified on the granulation tissues VWF+ collocation with RGS5, NOTCH3, and HEY1, as predicted by the maker gene plot (Figure 3F,G), confirming this new population of cells exist *in situ*.

**Figure 3.**
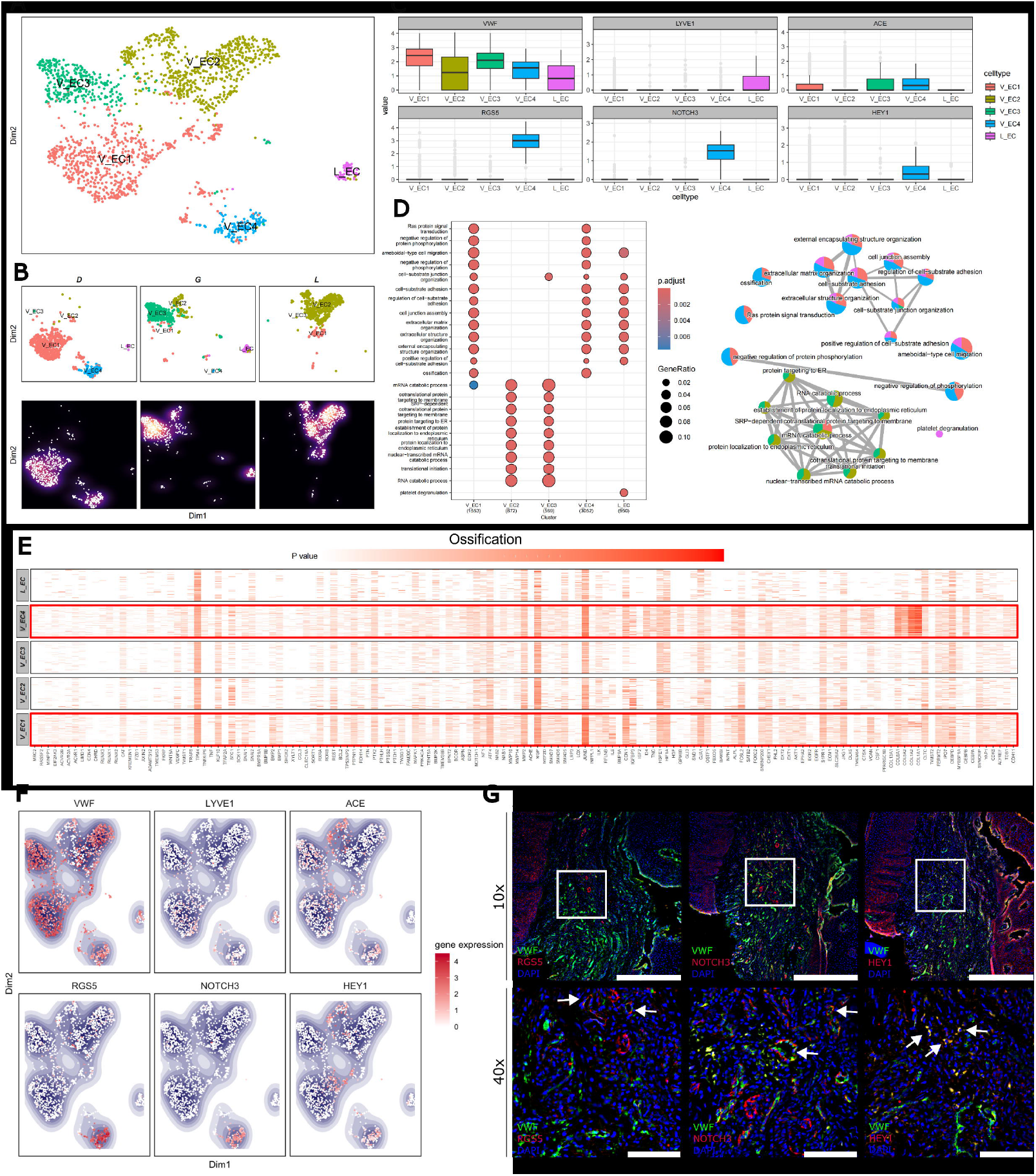
Endothelial subpopulation analysis. (A) Reclustering UMAP plot of Endothelial cells, single cells coloured by cluster annotation. (B) UMAP plot divided by tissue and population density plot. D refers to Diseased (Granulation tissue), G refers to Gingiva (Healthy), and L refers to Periodontal Ligament (Healthy). (C) *Box plot showing subset-specific markers and their expression levels within the endothelial sub-clusters*. (D) GO enrichment terms for comparing clusters, using the compareCluster function, and emapplot function from clusterProfiler. (E) *Ossification-enriched genes and their expression levels were plotted on a heatmap, where cluster VE_4 exhibited the highest expression levels of these genes. (F,G) Validations and mapping of unbiasedly identified populations based on the expression of selected marker genes. White arrows showing colocalisation; Scale bars: 10x 500μm, 40x 100μm*

To explore cluster-specific functions and track the developmental origin of cell types, pseudotime analysis was performed using Monocle2, and the starting point of differentiation was determined by CytoTRACE2 (Figure 4A-C). While lymphatic endothelial (L_EC) cells exhibited the lowest degree of differentiation, the granulation specific V_EC4 subpopulation, characterised by NOTCH3 expression and its involvement in ossification, demonstrates the highest degree of differentiation. Subsequently, we conducted GSEA comparing the disease tissue with gingiva and PDL (Figure 4D). The most significantly enriched pathways emphasize vascular development-related pathways.

**Figure 4.**
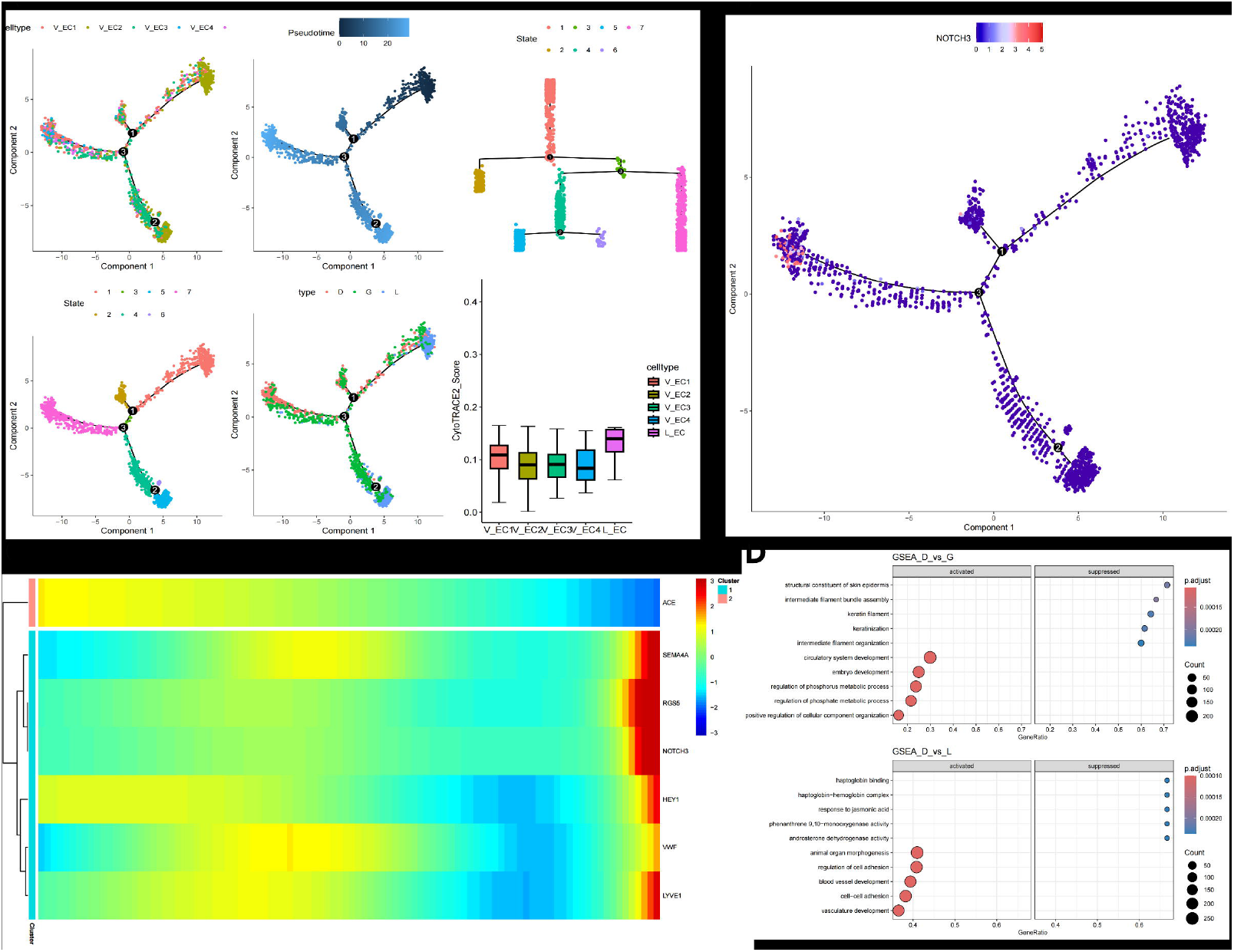
Pseudotime analysis elucidates the origin of endothelial cells in periodontal tissues. (A) Pseudotime plotting reveals the differentiation trajectory of endothelial cells, annotated by cell type, state, cluster, and tissue source. CytoTrace2 identifies the starting point of endothelial cell differentiation. (B) Pseudotime plotting showing high NOTCH3 specific expression where granulation-specific population V_EC4 is located. (C) Marker genes for endothelial cell identification are plotted along pseudotime, illustrating their dynamic expression profiles during differentiation. These genes are further clustered based on their expression patterns. (D) Gene set enrichment analysis (GSEA) identifies enriched pathways among different tissue resources. This comprehensive analysis provides insights into the developmental trajectory and functional characteristics of endothelial cells in periodontal tissues.

Given the potential functionality discovered of the endothelial compartment of the granulation tissue, we employed a cell-to-cell communication analysis to investigate the mechanism by which these populations regulate the environment. Our analysis revealed that the granulation tissue specific clusters (V_EC1 and V_EC4) displayed the biggest weight of interaction, with the V_EC4 cluster having the most robust signaling enrichment of all endothelial clusters(Figure 5A), suggesting the V_EC4 plays a crucial role in the diseased periodontal microenvironment. This population signaling is significantly marked by the NOTCH pathway and pathway interaction analysis revealed that V_EC4 interacts with VEGF, COLLAGEN, and BMP pathways, all crucial for vascular and ossification development (Figure 5B-F; Supplementary Figure 9). Our analysis suggests that the V_EC4 endothelial population is crucial for both vascular development and ossification of the microenvironment of periodontal disease.

**Figure 5.**
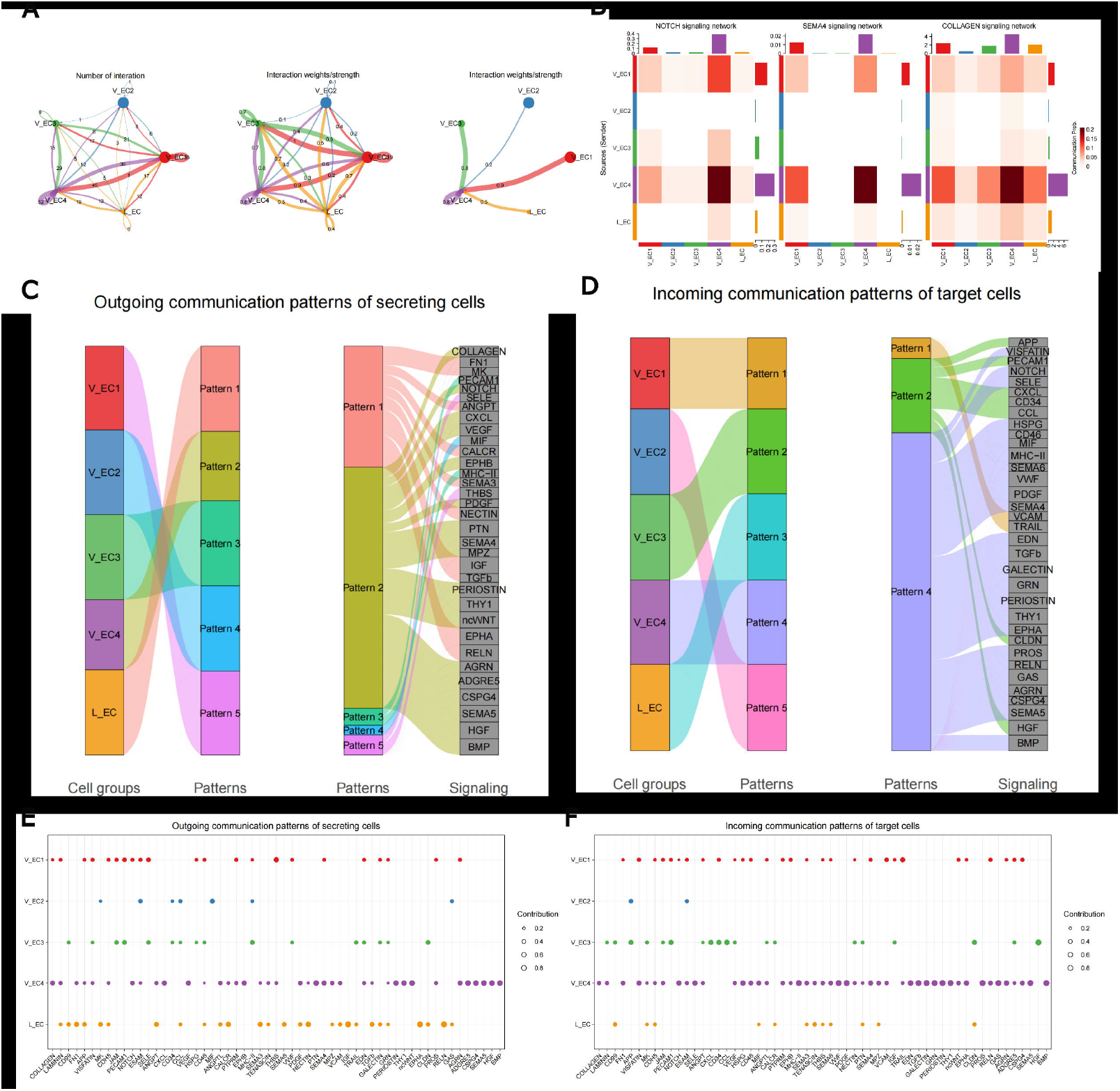
Cell-cell communication among endothelial types in the periodontal pathogenesis microenvironment. (A) Circos plots displaying the number and strength of cell-cell interactions among all endothelial cell sub-types across granulation tissue, ligament, and gingiva. Node size corresponds to the number of interactions, while edge width indicates the number of significant ligand-receptor pairs between two cell types. (B) Heatmaps summarizing specific signals between interacting cell types. Interactions are divided into outgoing and incoming events for specific cell types. The color gradient indicates the relative strength of the interactions. (C, D) Signaling sender and receiver information is presented on the riverplot and enriched as clusters. (E,F) Dot plot illustrates the altered signaling pathways among each subtype of endothelial cell. V_EC4 has enriched the most significant outgoing and incoming signaling.

## Discussion

In this study, we aimed to comprehensively elucidate the transcriptional landscape underlying periodontal granulation tissues through scRNA-seq analysis via comparing to the diseased tissue to healthy gingiva and periodontal ligament. Our findings shed light on the molecular complexity of these tissues, offering valuable insights into the pathogenesis of periodontal disease. Our experimental workflow enabled us to obtain high-resolution transcriptomic data from a diverse array of cell populations within the periodontal microenvironment, understanding the trajectory of differentiation from healthy to disease, as well as finding a novel endothelial cell population that could be the target of periodontal therapy.

Our scRNA-seq analysis unveiled a compelling insight into a distinct endothelial cell subpopulation (V_EC4) in granulation tissue, characterized by the NOTCH pathway signalling and visualised peripherally to blood vessels, a characteristic of pericytes. The endothelial derived NOTCH pathway expression is extensively documented to promote angiogenesis and osteogenesis in bone[22]. Additionally, NOTCH+ endothelial cells were co-localised to RGS5 and HEY1. RGS5 has been described in the dental pulp as a source of stem cells to produce odontoblasts[23]. Hey1 is a downstream effector of the canonical Notch signalling[24], and the TGF-b/BMP signalling, independently of Notch[25]. Our results suggest that this population plays a crucial potential role in shaping the diseased microenvironment, emphasizing the close association between vascular development and ossification processes. Further functional analysis is needed to confirm these results, however targeting this population would be a potential novel therapeutic target for enhancing periodontal regeneration.

Moreover, our findings underscore the dynamic interplay between various cell types within the periodontal disease progression-related tissue, highlighting that Fibroblasts are the most interactive cell type during disease progression. Fibroblasts, particularly the D_FB1 subcluster, exhibit a progenitor-like state with pronounced extracellular matrix (ECM) organization activity. This population’s low differentiation state and its association with the Wnt signaling pathway suggest a crucial role in the scar-like fibrotic phenotype observed in periodontal disease. A biological similar event takes place with Cancer-associated fibroblasts (CAFs), where they have potency to differentiate into a functional fibroblast that produce ECM structures and metabolic and immune reprogramming of the tumour microenvironment impacting tumour progression[26]. Therefore, further understanding of how to modulate the fibroblast expression and phenotype in the granulation tissue could be crucial for managing the progression of periodontal disease.

Clinically, there is debate as to whether the granulation issues should be left or removed. Our dataset shows that the microenvironment of the granulation tissue is intricate, with populations supporting removal of the tissue and populations suggesting maintaining it. What is certain is that current clinically available materials used in periodontal disease management are not based on the biological pathways presented in our research[27]. Therefore, there is scope for developing new biomaterials that stimulate or repress genes crucial for periodontal disease progression or that induce regeneration, advancing therefore periodontal disease management.

In conclusion, the study advances the current understanding of the pathogenesis of periodontal disease by characterising the most important regulatory compartments of the granulation tissue and the understanding their developmental origins in relation to periodontal healthy tissues.

## AUTHOR CONTRIBUTIONS

VCMN conceptualised, designed, and performed experiments; curated and analysed the data; and wrote, edited, and reviewed the original manuscript. WZ performed the bioinformatic analysis, wrote, edited, and reviewed the original manuscript. KF and PD performed the single cell RNAseq. JK critically revised the manuscript. CZ, PS, and LN provided resources and reviewed the manuscript. All co-authors reviewed the manuscript.

## ACKNOWLEDGEMENTS

We thank all the patients who contributed to this study, the support of the Periodontology MClinDent students and GSTT nursing staff at Guy’s Hospital.

## DATA AVAILABILITY

All data associated with this study are present in the paper or the Supplementary Materials. Requests for data should be addressed to the corresponding authors.

## Supplementary Figures Legend

**Figure S1. Overview of single cells from granulation tissue in periodontitis (PD)** (A) Schematic overview of the study. (B) Uniform Manifold Approximation and Projection (UMAP) representation showing nine cell types identified in the granulation tissue. (C) Selected markers used for cell type identification. Notably, FCGR3D and ntt55 CSF3R, typically associated with neutrophil markers, were observed; however, the cell number was insufficient to form a distinct cluster. Differentially expressed genes (DEGs) were normalized and visualized on a heatmap, displaying the Top 10 genes ranked by log2 fold change (log2FC). (D) UMAP from (B), colored by the tissue origin of patients. (E) Hematoxylin and eosin (HE) staining of biopsies from granulation tissue.

**Figure S2. KEGG Pathway Enrichment Analysis of Granulation Tissue Cell Types** Within the granulation tissue, KEGG pathways enriched using and shown by dotplot from the function of clusterProfiler, categorized by pathway type and split by cell types. Pathways are ranked by adjusted p-value (P.adjust), with a significance threshold set at pvalueCutoff = 0.05.

**Figure S3. Cell-Cell Interaction Dynamics Across Tissues** (A) Circos plots displaying the number and strength of cell-cell interactions among all cell types across granulation tissue, ligament, and gingiva. Total incoming and outgoing signaling strengths are illustrated for each cell type. Fibroblasts exhibit stronger interaction as receivers than as senders, while endothelial cells demonstrate stronger interaction as senders than as receivers. (B) Heatmaps summarizing specific signals between interacting cell types. Interactions are divided into outgoing and incoming events for specific cell types. The color gradient indicates the relative strength of the interactions.

**Figure S4. Reclustering and Functional Analysis of Epithelial Cells** (A) UMAP plot showing reclustering of epithelial cells, with individual cells colored by cluster annotation. (B) UMAP representation of epithelial cells, differentiated by tissue origin, accompanied by density plots for each tissue. (C) Heatmap displaying differentially expressed genes (DEGs) among epithelial cell subclusters, ranked by log2 fold change (log2FC). (D) Gene Ontology (GO) enrichment analysis comparing functional differences across epithelial cell clusters, performed using the compareCluster function. (E) Visualization of GO enrichment results using the emapplot function from clusterProfiler, illustrating how each cluster contributes to enriched pathways. (F) Rate of cell annotation from each tissue resource. (G) Trajectory analysis revealing differentiation relationships among epithelial cells. Differentiation starting points were identified using CytoTRACE2, and pseudotime trajectories were inferred using Monocle2. Trajectory plots were visualised by pseudo time, tissue origin, cell type and differentiation stage.

**Figure S5. Relative Expression of Epithelial Cell Markers** Bar plots showing the relative expression levels of typical epithelial cell markers across each cluster.

**Figure S6. Reclustering and Functional Analysis of Mesenchymal Stem Cells (MSCs)** (A) UMAP plot showing the reclustering of Mesenchymal Stem Cells (MSCs), with individual cells colored by cluster annotation. (B) UMAP representation of MSCs differentiated by tissue origin, accompanied by density plots for each tissue. (C) Heatmap illustrating differentially expressed genes (DEGs) among MSC subclusters, ranked by log2 fold change (log2FC). (D) Gene Ontology (GO) enrichment analysis comparing functional differences across MSC clusters, performed using the compareCluster function. (E) Visualization of GO enrichment results using the emapplot function from clusterProfiler, demonstrating contributions of each cluster to the enriched pathways. (F) Rate of cell annotation from each tissue resource. (G) Trajectory analysis revealing differentiation relationships among MSCs. Differentiation starting points were identified using CytoTRACE2, and pseudotime trajectories were inferred using Monocle2. Trajectory plots are displayed based on pseudotime, tissue origin, cell type, and differentiation stage.

**Figure S7**. Bar plots showing the LEF1 expression levels on different fibroblast clusters

**Figure S8**. Bar plots showing the relative expression levels of typical fibroblast cell markers across each cluster.

**Figure S9**. The dot plot illustrates all significant ligand-receptor pairs contributing to signaling among endothelial subclusters in the dataset. The color of each dot represents the calculated communication probability, while the size of the dot corresponds to the associated p-value. Empty spaces indicate instances where the communication probability is zero. P-values were computed using a one-sided permutation test.

